# Mosquito Larvae (*Culex spp.*) Startle Responses to Vibration Stimuli

**DOI:** 10.1101/2021.02.18.431787

**Authors:** Ellie R. Harrison, William J. Budenberg

## Abstract

We explored startle responses of *Culex spp.* larvae to varying frequencies and durations of vibrations, before investigating habituation (a behavioural learning phenomenon). The startle response consists of a rapid movement to permit escape from a potentially dangerous stimulus. We used the automated Zantiks MWP_vb unit to analyse larval startle responses through the built-in live video tracking. Set within the unit’s controlled environmental chamber, we recorded distance travelled by the larvae. These results were used to optimise the vibration startle response for use in habituation studies. Repeated vibrations appeared to reduce larvae startle responses, although further analysis is required to confirm habituation. This study not only provides an insight into the learning ability of mosquito larvae, but demonstrates the ease, replicability and control conferred by the automated unit. Furthermore, this method is applicable to many other organisms and shows potential for more sophisticated environmental effect testing, such as behavioural toxicology.

## Introduction

The ability to respond to potentially dangerous stimuli is fundamental to evolutionary success. Startle responses involve a rapid response to a sudden stimulus and therefore facilitate escape from a threatening situation. These escape responses function to terminate a negative experience or hazard exposure (Patterson-Kane, 2018) and are species, age and context specific. As a significant contributor to a species’ survival, escape responses can often be reliably produced, thus permitting research into the underlying mechanisms of these behaviours.

Performing stereotypical escape responses (Baglan et al., 2017), aquatic mosquito larvae are excellent animals for further study. Surface-dwelling larvae rapidly dive with the onset of danger, before recovery to the water’s surface in the absence of stress. Yet repeated diving has been shown to increase larval mortality (Tuno et al., 2004) and impact development (Lucas & Romoser, 2001) due to the significant energy expenditure. The resulting trade-off between escaping potential danger versus wastage of energy is of evolutionary significance to the mosquitoes. One such solution is habituation, a widely characterised phenomenon involving a progressive and reversible decrease in a behavioural response resulting from a repeated stimulus (Rankin et al., 2009). Habituation must be distinguished from sensory adaptation, whereby the sensitivity of receptors is lost, as well as effector fatigue, involving response system depletion. Moreover, habituation is stimulus specific, thus introduction of a new stimulus modality will still elicit a response. Although studied in numerous model organisms, the investigation of habituation in mosquitoes is relatively novel. Habituation to a visual stimulus has been demonstrated in *Aedes aegypti* larvae (Baglan et al., 2017).

This study aimed to investigate habituation of *Culex spp.* larvae to a vibration stimulus mimicking predator-approach. Using the Zantiks MWP unit automated and controlled environmental setup, we characterised the optimum frequency and duration of vibration to elicit the escape response. These results were used to inform experiments investigating habituation.

## Materials and Methods

### Animals

The experiments used fourth-instar larvae of *Culex spp*. collected from a local wild reservoir. Mosquitoes were reared in environmental day-night cycles and temperatures. Genus identification was achieved via microscopy (**Fig.1**) and unique behaviour; however, the sex and species of the larvae is unknown. Each experiment ran one or more distinct 24-well plates of larvae. After experimentation, the larvae were returned to a separate local reservoir.

**Fig.1.**
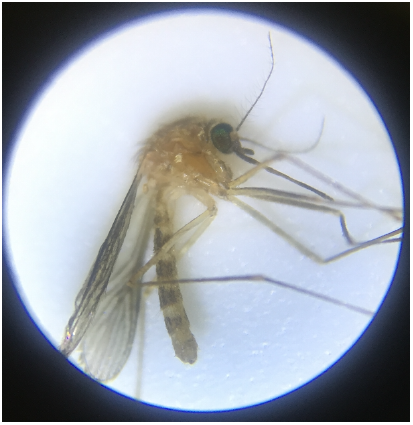
microscopy image for confirmation of genus *Culex spp*.

### Experimental Protocol

Environmental control and procedure were achieved within the Zantiks MWP_vb unit (Zantiks Ltd, Cambridge *https://zantiks.com/*). The unit maintained a constant white 500 lux overhead light and a temperature of 20°C, consistent with the average environmental temperature of late summer in the UK. Each larva was placed in a well within a 24-well plate, filled with the reservoir’s water. The 24-well plate was inserted into the unit, which reported the distance travelled by the larvae second-by-second using the integrated video tracking (**Video.S1**: available online). Plates were secured with sticky tack to prevent gross movement of the plate as a result of the vibration stimulus. Each experiment included a 30-minute acclimation time to allow the familiarisation of larvae to the new environment. 5-minutes between startles allowed larvae recovery (2.5-mins Pre and Post) (**Fig.2**). ‘TYPE’ refers to vibration ‘Startle’ or no vibration ‘Pre’. Due to the use of two units, we also investigated whether responses differed between units through the ‘TYPE:UNIT’ interaction.

**Fig.2.**
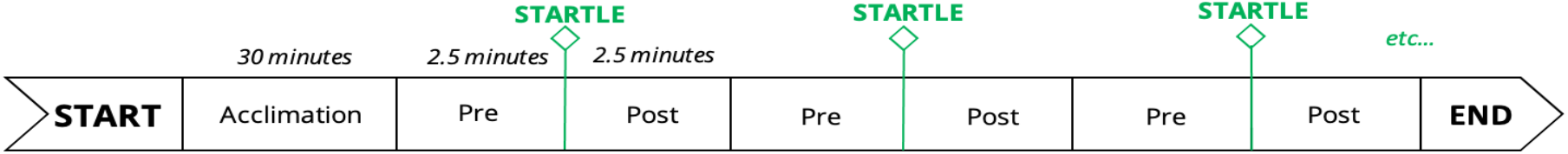
diagram showing the base experimental protocol within the Zantiks MWP unit.

### Vibration frequencies and durations

The aim of the experiment was to establish the frequency resulting in the greatest distance travelled. The inbuilt motor of the unit vibrates the Zantiks plate, thus coding of the motor’s parameters controls the characteristics of the vibration. Firstly, two 24-well plates of larvae were run with frequencies from 58Hz, increasing by a factor of 2, up to 1860Hz in two separate units. Plates were swapped between units before running the experiment again. These results informed the narrowing of the frequency range. Another experiment with two plates of fresh larvae was carried out, increasing the frequency at 1.5x increments from 232Hz to 1174Hz. Plates were swapped between units and the protocol repeated.

The same principles were applied to duration experiments. Using the optimal frequency previously determined, two 24-well plates of larvae were subject to vibrations ascending from 2ms to 550ms. After swapping the plates between units, another repetition was carried out. Subsequently, a narrow range duration experiment was undertaken, with doubling increments from 70ms to 550ms. Plates were swapped and repeated.

### Habituation

Three experimental protocols aimed to investigate whether habituation to vibrations occurred in wild *Culex spp.* larvae. The first protocol replicated the experimental design of Baglan et al., 2017, consisting of 15 stimuli separated by 5 minutes. All vibrations were set at the optimum frequency and lasted 0.5s. Two plates were run in separate units with one repetition. The subsequent experiment mirrored the first protocol but reduced the inter-stimulus time to 2 minutes. The final experiment consisted of 15 consecutive stimuli separated by 10 seconds, repeated in 4 blocks separated by 5 minutes. A ‘Test’ stimulus 5 minutes after each block was included to investigate the presence of habituation retention.

### Statistical tests

We performed statistical analyses with “R” software, version 4.0.2 (R Core Team, 2013). Type III, unbalanced ANOVA analyses were used to test for significance between no vibration and vibration. Additionally, pairwise T test comparisons with Bonferroni correction were used to compare different vibration durations. Due to the nature of our data and the large number of second-by-second samples, typical tests of normality were inappropriate. Upon visualisation of the data, it was apparent that there were a large number of ‘0’ entries where larvae had not moved. As a result of this positive skew, normality cannot be assumed. Nevertheless, our large sample size - and hence statistical power - combined with the robustness of ANOVA to violations of normality (Blanca et al., 2017), suggest our results are valid.

## Results

Firstly, we aimed to establish the optimum frequency for eliciting the startle response in the larvae. A preliminary wide range frequency experiment, with frequencies from 58Hz to 1856Hz, allowed identification of a narrower range to investigate further. The second experiments consisted of frequencies increasing from 232Hz to 1174Hz. **Fig.3A** shows the boxplot of the different frequencies against distance. No clear trend is visible, except a notable drop at 522Hz which is consistent in both units (**Fig.3B**).

**Fig.3.**
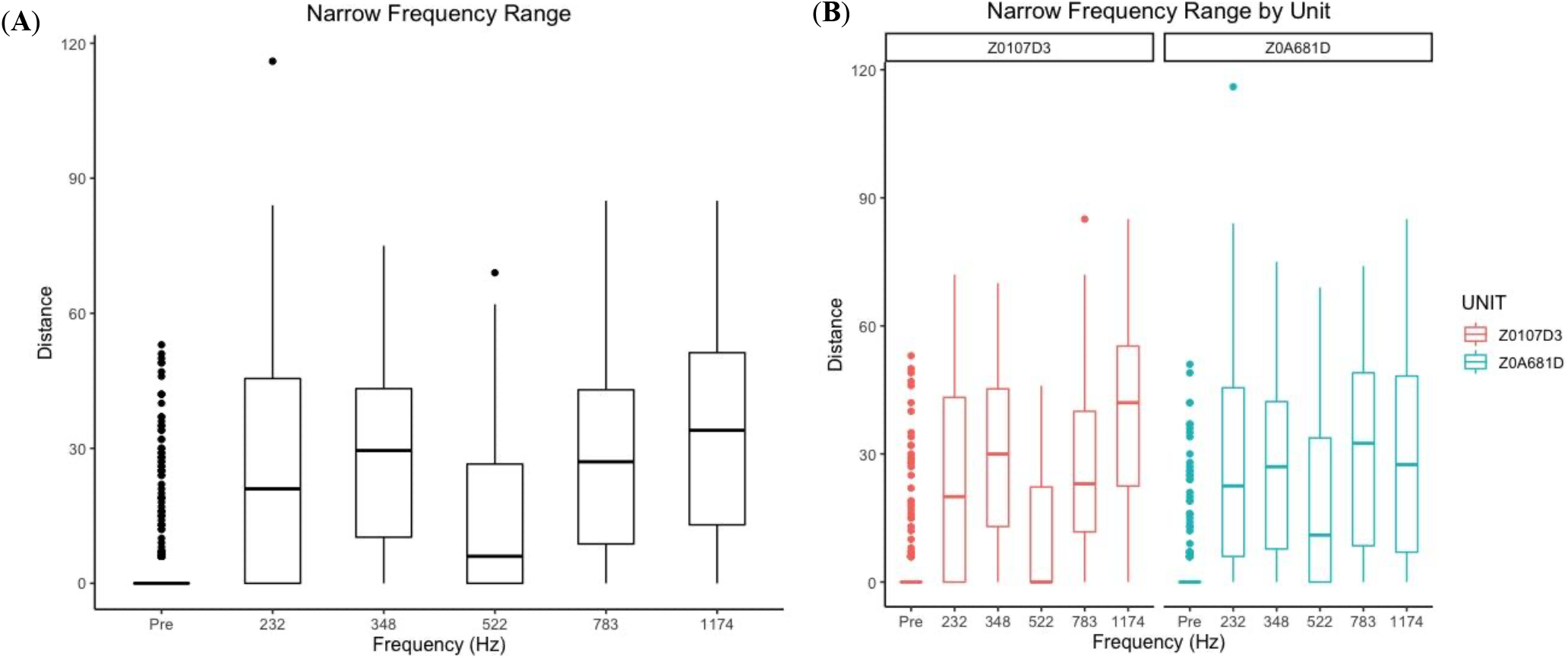
**(A)** Narrow frequency range box plot comparing frequency to distance travelled by the larvae. All the vibrations are greater than the no vibration Pre. However, 522Hz shows a marked decrease compared to the other frequencies. **(B)** Narrow frequency range box plot split by unit. Both units show the decrease at 522Hz, indicating a consistent issue within the units.

This result may seem anomalous but could be explained by the vibration transfer from the motor through other components, such as the diffusing stand. The differing harmonic frequencies of these components modulate the motor generated signal, resulting in the larva receiving an altered vibration frequency. Nevertheless, the signal received is consistent across repeated stimuli and different units.

ANOVA analysis (**Table 1**) produced an extremely significant effect of ‘TYPE’ (i.e. vibration versus ‘Pre’), thus confirming the vibrations produce a significant response from the Pre control. Additionally, the significant ‘TYPE:UNIT’ interaction could be explained by the unit harmonics issue. From this exploratory experiment, 900Hz was decided upon as an optimal frequency for producing a strong escape response.

**Table 1.** Type III ANOVA table for the narrow frequency range. TYPE is highly significant (i.e. vibrations are significant from no vibration Pre) and there is a significant interaction between TYPE and UNIT. F and Pr(>F) values 3.s.f. Signif. codes: 0 = *** 0.001 = **

| Narrow Frequency ANOVA |  |  |  |  |
| --- | --- | --- | --- | --- |
|  | Sum Sq | DF | F value | Pr(>F) |
| Intercept | 3534 | 1 | 12.7 | 0.000391 *** |
| **TYPE** | **75577** | **5** | **54.2** | **< 2.2e-16 \*\*\*** |
| UNIT | 2 | 1 | 0.0059 | 0.939 |
| **TYPE:UNIT** | **4381** | **5** | **3.14** | **0.00810 \*\*** |
| Residuals | 264552 | 948 |  |  |

Two similar experiments were carried out looking into the effect of vibration duration on distance travelled using vibrations of 900Hz of varying lengths. A first experiment, using durations from 2ms to 550ms, was utilised to narrow down the range for the second experiment. **Fig.4** depicts the boxplot of the vibration durations against distance travelled, with vibrations doubling from 69ms to 550ms. ANOVA analysis revealed a very strong significance of ‘TYPE’ but no significant interaction with ‘UNIT’ (**Table 2**), confirming only vibrations had a significant effect on distance travelled. We compared different durations with pairwise T test comparisons. The first significant comparison is between the durations of 137ms to 275ms, from no startle to startle, so we can conclude 900Hz vibrations lasting 275ms plus will elicit the escape response. Furthermore, the comparisons highlight the greatest startling occurs at 550ms, thus we decided to use vibrations of 900Hz lasting 550ms in the subsequent habituation experiment.

**Fig.4.**
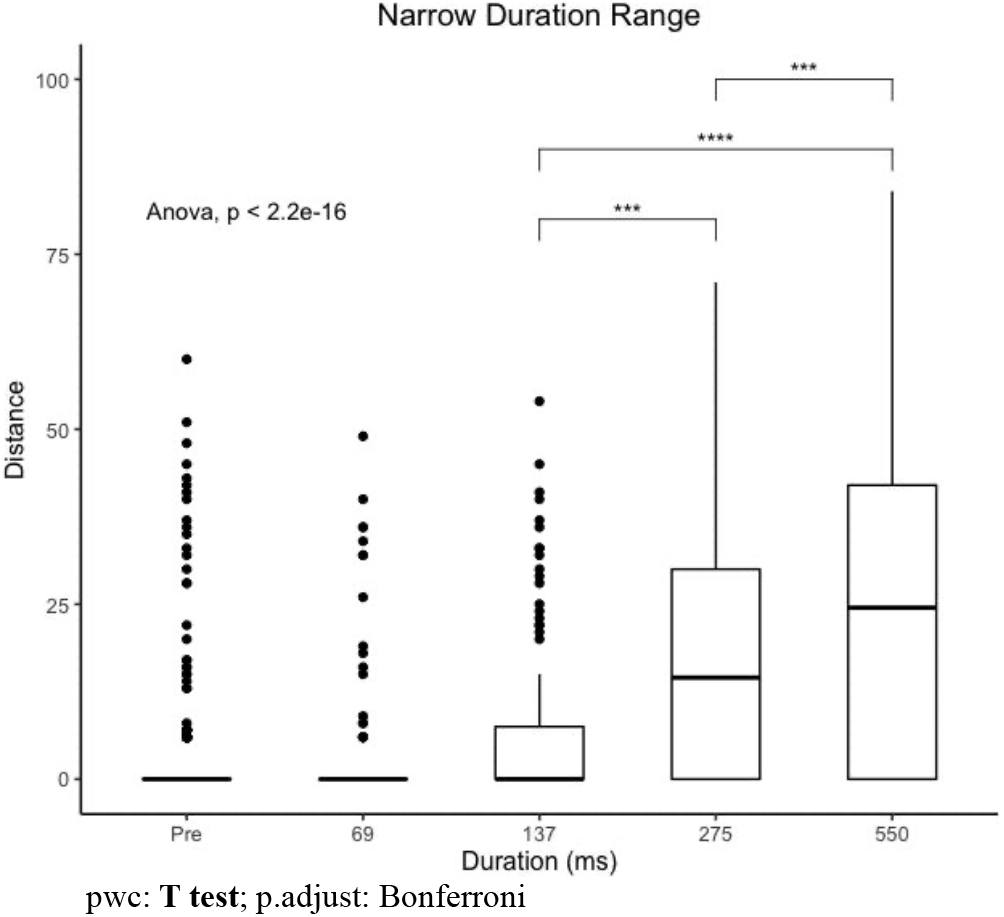
Narrow duration boxplot comparing duration to distance travelled. There is little response until 137ms+. The significant pairwise T-tests show a response elicited from 275ms and an even greater response with 550ms.

**Table 2.** Type III ANOVA table for the narrow duration range. TYPE is highly significant (i.e. vibrations are significant from no vibration Pre). There is no significant interaction between TYPE and UNIT indicating consistent results across the two units. F and Pr(>F) values 3.s.f. Signif. codes: 0 = *** 0. 0.001 = **

| Narrow Duration ANOVA |  |  |  |  |
| --- | --- | --- | --- | --- |
|  | Sum Sq | DF | F value | Pr(>F) |
| Intercept | 1813 | 1 | 10.5 | 0.00123 ** |
| **TYPE** | **32472** | **4** | **54.2** | **< 2e-16 \*\*\*** |
| UNIT | 59 | 1 | 47.1 | 0.557 |
| TYPE:UNIT | 1165 | 4 | 1.69 | 0.150 |
| Residuals | 130592 | 758 |  |  |

The final experiment aimed to investigate habituation of the startle response to the vibrations. If habituation were occurring, we would expect to see a progressive decrease in the distance travelled by the larvae as the number of vibration trials increases. After varying the length of interstimulus time, 10-seconds was decided as the optimum value to allow larvae recovery to their surface-dwelling position. The experiment involved 4 blocks of 15 trials of startles separated by 10-seconds, with 5 minutes between blocks. **Fig.5** shows the 15 startles repeated 4 times. A ‘Test’ startle was carried out 5 minutes after the 4 blocks to investigate effector fatigue (response system depletion). For blocks 1-3, we used the first trial of the subsequent block as a ‘Test’ startle.

**Fig.5.**
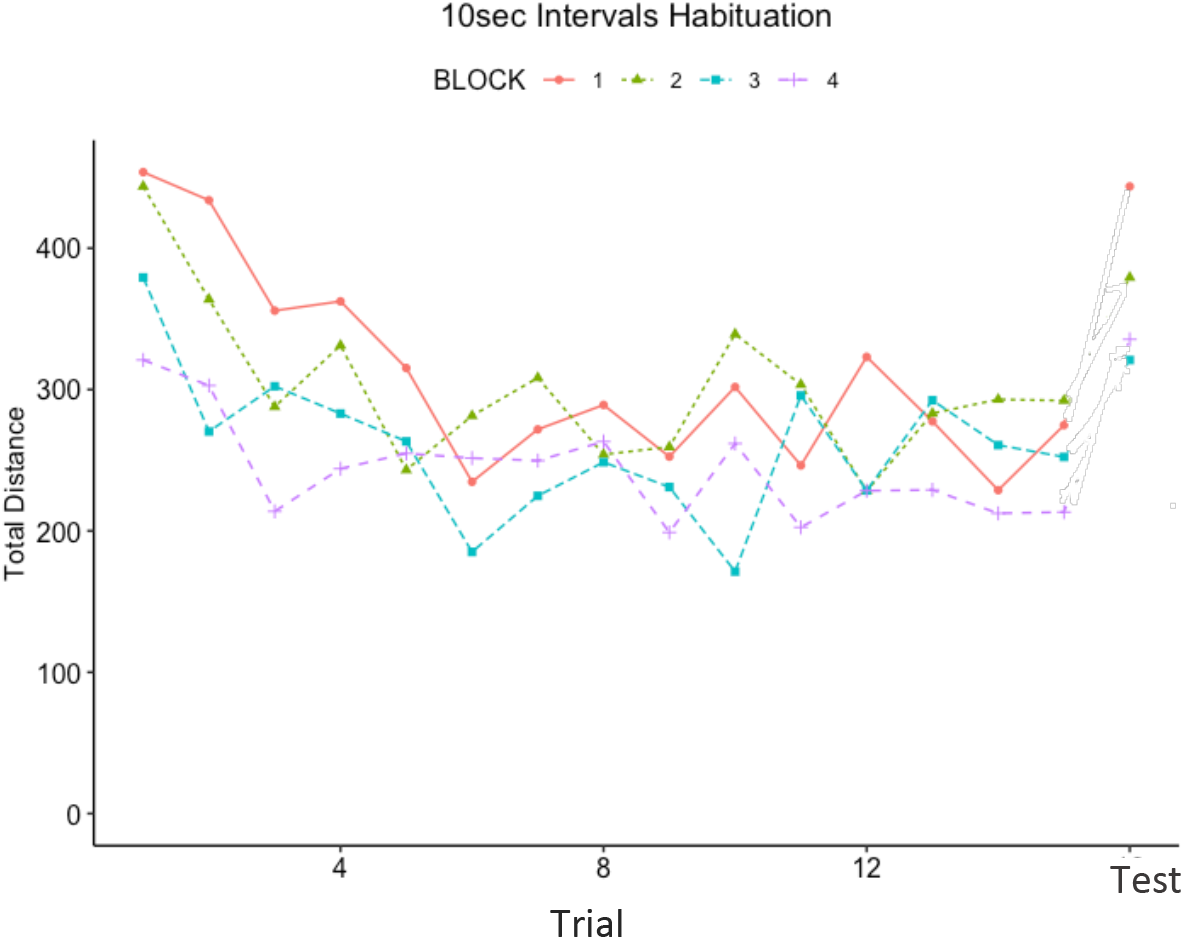
Habituation graph plotting total distance of the larvae against time with 10-second intervals between trials. Each 15-trial vibration block is represented by a different colour. The ‘Test’ startle for each block occurred 5-minutes after the 15-trials. For blocks 1-3, the ‘Test’ startle is the first trial of the subsequent block e.g. trial 1 of block 2 is the ‘Test’ startle for block 1.

There appears to be a downward trend in startling for all 4 blocks over the course of the 15 trials, but the decrease is only ~30%. Additionally, the ‘Test’ startle for each block elicits a greater distance travelled compared to the 15^th^ trial but does not recover fully to the starting 1^st^ trial distance. Effector fatigue over the course of the experiment is a likely explanation, but it may also be explained by long-term learning of the stimulus. The previous work with *Aedes aegypti* larvae suggests a 1-hour long mnesic trace to the visual stimulus was established (Baglan et al., 2017). Although not conclusive, this experiment does indicate the possibility of habituation of the *Culex spp.* larvae to vibration stimuli.

## Discussion

This project successfully characterised vibration frequencies and durations to elicit startling in *Culex spp.* larvae. For maximal distance travelled during the startle, a vibration with a frequency between 700-1200Hz lasting 550ms should be used. These cut-off values can inform pre-pulse inhibition (PPI) experiments, whereby it is possible to control whether a vibration results in startling. PPI describes a weaker pre-pulse stimulus, to which there is no startle response, inhibiting the startle reaction to a stronger stimulus. Therefore, these results permit an informed choice of the frequency and duration parameters to use for the ‘pre-pulse’ and ‘pulse’ vibrations.

The investigation into habituation of the *Culex spp.* larvae was inconclusive. We observed a downward trend in startle distance over the 15 trials, but the decrease was not confirmed as habituation. However, it is clear that we were unable to replicate Baglan *et al*’s habituation with an inter-stimulus length of 5 minutes (Baglan et al., 2017). Instead, under our experimental conditions, a smaller inter-stimulus length was required to observe learning. To rule out sensory adaptation, a different stimulus of the same modality is necessary. Additionally, effector fatigue can be further investigated via the inclusion of an alternative stimulus modality, such as a visual stimulus. If escape responses are elicited to both of these stimuli, there is a greater confidence that habituation has occurred.

Overall, the standardisation possible with the Zantiks units means this method has great potential for more sophisticated environmental effect testing on a wide range of organisms. One example is ecotoxicology, where this method allows quantification of the effect of chemicals on organisms’ behaviour. For example, the effect of the herbicide glyphosate on habituation has been characterised for 20 *Aedes aegypti* larvae (Baglan, Lazzari, & Guerrieri, 2018). Use of our automated method would permit a significant increase in sample size and thus statistical power.

With the mosquito’s role as a disease vector set to increase with global warming (European Commission Joint Research Centre, 2018), characterising the behavioural and learning mechanisms of this species is important. The increased behavioural understanding may underpin the development of effective interventions.

## End Matter

### Author Contributions and Notes

ERH joined Zantiks as a part time intern to carry out this research.

WJB & ERH designed the study jointly. ERH carried out most of the experiments, all the data analysis and wrote the paper.

WJB is the founder of Zantiks, and is such an interested party in the use of the Zantiks units in behaviour studies.

## Acknowledgments

This study was entirely funded internally by Zantiks as part of its research and development program.

